# Hypoxic Training Increases the Concentration of Serum Irisin and Reduces Weight in Diet-induced Obese Rats

**DOI:** 10.1101/500603

**Authors:** Zhigang Li, Xiquan Weng, Xue Song, Fangfang Zhao, Hao Chen, Wengtao Lin

## Abstract

Irisin promotes browning of white fat, improves energy metabolism, and-weight loss. In this study, we investigated the effects of different oxygen concentrations during hypoxic training on the serum irisin and the PGC-1α(peroxisome proliferator-activated receptor gamma coactivator 1-alpha)-FNDC5(fibronectin type III domain containing 5)-UCPl(uncoupling protein 1) signaling pathway in the skeletal muscle of obese rats. Male Sprague-Dawley Obese rats (n=80) were randomly divided into 8 groups as follows: the control group (group A, n=10); the endurance exercise group (AE group, n=10), which involved animal treadmill training at slope 0°, 20 m/min, 40 min/d, and 5 d/w; the 16.3% hypoxia exposure group (group B, n=10), 13.3% hypoxia exposure group (group C, n=10), and 11.3% hypoxia exposure group (group D, n=10), which were exposed to a low oxygen environment with oxygen concentrations of 16.3%, 13.3%, and 11.3%, respectively, for 12 h/d; and the 16.3% hypoxic training group (BE group, n=10), 13.3% hypoxic training group (CE group, n=10), and 11.3% hypoxic training group (DE group, n=10) with animal treadmill training during hypoxia exposure. After 8 weeks, the serum irisin concentrations in the AE, BE, CE, and DE groups were significantly higher than that in the A group (p<0.05). Hypoxia exposure and hypoxic training at the three different concentrations significantly increased PGC-1α and FNDC5 gene expression in the skeletal muscle. The PGC-1α and FNDC5 protein contents were significantly higher in the skeletal muscle of the obese rats in the C, AE, and DE groups than those in group A (p<0.05). UCP1 protein expression was significantly higher in groups C, CE, D, and DE than in group A (p<0.05). To conclude, training at oxygen concentrations of 13.3% and 11.3% significantly increased the serum irisin level, and 11.3% hypoxic training enhanced the effects of the PGC-lα-Irisin-UCP1 signaling pathway in skeletal muscle.

## Introduction

Obesity has become a global public health problem, and the rates of obese and overweight individuals of different genders and age groups continue to increase[1]. Therefore, safe weight-control methods and effective weight-loss action targets are a focus of research. As a new myokine[2] and adipokine[3], irisin causes browning of white fat and promote energy metabolism. Thus, irisin has great potential for the prevention and treatment of metabolic diseases and obesity. Numerous studies have found that exercise can stimulate skeletal muscle to release irisin into the blood circulation and that it has obvious benign effects on obese or diabetic patients of different ages[4–8]. Currently, the relationship between irisin and various chronic diseases related to energy metabolism disorders is gradually being revealed, and new functions of irisin are constantly being discovered[9, 10]. Long-term endurance training or short-term sprint training can both promote an increase in the irisin level[11–13]. However, at present, how irisin regulates energy metabolism is not clear, and there has been no agreement on the effects of different exercise methods on the irisin concentration in human blood circulation.

Hypoxic stimulation can increase the basal metabolic rate of the human body[14], enhance energy output, and reduce appetite[15]. Hypoxic intervention has a significant effect on improving obesity and related metabolic syndromes[16,17]. Hypoxic training is a superposition of hypoxia and exercise stimulus. An appropriate hypoxic training intervention can achieve an ideal weight-control effect[18]that is more effective than hypoxia or an exercise intervention alone[19, 20]. However, in weight loss schemes involving hypoxic training, the oxygen concentration, mode of hypoxia, and exercise regimen are not the same, and the results have been controversial[21]. The low oxygen concentration in animal experiments is not suitable for humans. Therefore, hypoxic training for the purpose of weight loss needs further exploration. In addition, few studies have investigated the effects of hypoxic training on irisin, and whether weight loss can be achieved by promoting irisin release is unclear. Based on these facts, an intervention with intermittent hypoxia exposure to three low-oxygen concentrations combined with endurance training was employed in this study to explore the appropriate stimulating oxygen concentration for hypoxic training-assisted weight loss, to analyze the stimulation effects of hypoxic training on irisin, and to enrich the theory concerning exercise weight loss.

## Materials and methods

### Experimental animals

A total of 140 male Sprague-Dawley rats aged 6 weeks that were specific pathogen-free (SPF) grade and weighed 194.70±1.24 g after one week of adaptive feeding were purchased from the Experimental Animal Center of Southern Medical University (license number: SCXX (Yue) 2011-0015). The rats were housed in cages (4 per cage) with an ambient temperature of 25-26°C, a relative humidity of 45% to 55%, natural light, and a freely available diet. The daily food intake of the rats was recorded, and the body weights were recorded once weekly.

### Ethics Statement

All animal experiments were carried out in accordance with the National Institutes of Health Guide for the Care and Use of Laboratory Animals and were approved by Ethics Committee of Guangzhou Sport University (approval No:2016DWLL-001). All surgery was performed under appropriate anesthesia, and all efforts were made to minimize suffering.

### Diet and exercise intervention

The rats were randomly divided into the normal diet group (N group, n=20) and the high-fat diet group (HFD group, n=120), which was fed a HFD. No significant difference was observed in the initial average body weights between the two groups (p>0.05). The feed was provided by the Guangdong Medical Laboratory Animal Center. The high-fat animal model feed was composed of a total energy of 4.5 kcal/g, a mass ratio of 19% protein, 18.5% fat, and 50.5% carbohydrate and had a caloric ratio of 17.5% protein, 37% fat, and 45.5% carbohydrate.

After 8 weeks of feeding, 16 rats in the HFD group and 10 in the N group were randomly selected. The body weights and body lengths were measured, and Lee’s index was calculated (calculation formula: body weight (g)^1/3^ / body length (cm) × 10^3^). Orbital blood was collected; the blood glucose (BG) concentration was measured using GLUCOCARD^TM^ Test Strip II (ARKRAY, Japan), and the blood lipid indexes were measured after separating the sera. An average body weight exceeding 20% of that of the N group rats and significant differences in Lee’s index, BG, and the blood lipid indexes were used as standards[22, 23]to verify the success of nutritionally obese rat model construction.

Eighty eligible obese rats were randomly divided into 8 groups as follows: the normal oxygen quiet group (group A, n=10) without any intervention; the aerobic exercise group (AE group, n=10) with endurance training under a normal oxygen environment; the 16.3% oxygen concentration quiet group (group B, n=10); the 16.3% oxygen concentration exercise group (BE group, n=10); the 13.3% oxygen concentration quiet group (C group, n=10); the 13.3% oxygen concentration exercise group (CE group, n=10); the 11.3% oxygen concentration quiet group (D group, n=10); and the 11.3% oxygen concentration exercise group (DE group, n=10). No significant differences was observed in the average weights of the rats in the 8 groups (p>0.05). Rats in groups B, C, and D were exposed to a low oxygen environment with oxygen concentrations of 16.3%, 13.3%, and 11.3%, respectively, for 12 h/d. Hypoxia exposure was administered with a low oxygen partial pressure system (HTS) manufactured by Hypoxico Co., USA. The hypoxia exposure interventions in the BE, CE, and DE groups were the same as those in groups B, C, and D, and moderate-intensity treadmill training was performed during hypoxia exposure[24]. The exercise regimen was a 0° slope, 20 m/min, 40 min/d, and 5 d/w. The intervention period was 8 weeks following 2 weeks of adaptive training before the formal exercise intervention. During the intervention, the rats were fed the HFD continuously.

### Animal sampling

After the intervention, the body weights and body lengths of the rats were measured after fasting for 12 h. Anesthesia was performed by intraperitoneal injection of 10% chloral hydrate (0.3 mL/100 g). Blood was taken from the abdominal aorta, and the blood collection tubes were allowed to stand at room temperature for 30 min. The blood was centrifuged at 4°C and 3000 r/min for 15 min, and the supernatant was collected and quickly placed in a −80°C freezer for testing. Mesenteric, bilateral perirenal, epididymal, and groin fat and the bilateral soleus, gastrocnemius, and quadriceps muscles were collected, washed with 0.9% saline, weighed, frozen in liquid nitrogen, and stored at −80°C.

### Enzyme-linked immunosorbent assay (ELISA)

The serum irisin concentration were measured using the Irisin Competitive ELISA kit (AdipoGen, LOT: K1451507,Switierland), the adiponectin levels were measured using the (rat) ELISA kit (AdipoGen, LOT: K2371605, Switierland), leptin was measured using the Rat Leptin ELISA Kit (Millipore, LOT: 2851208,USA), and the insulin levels were measured using the Rat/Mouse Insulin ELISA Kit (Millipore, LOT: 2811046,USA). The ELISA test procedures were carried out in strict accordance with the manufacturer’s instructions. Each sample was subjected to duplicate detection. A multiband full-wavelength scanner (Thermo Fisher Scientific, USA) was used in the assays. The concentrations of total cholesterol (TC), triacylglycerol (TG), low-density lipoprotein cholesterol(LDL-C), and high density lipoprotein cholesterol (HDL-C) in the serum were measured by enzymatic colorimetric assays using commercially available detection kits (Biosino Biotechnology Co, Ltd, Beijing, China). The homeostasis model assessment of insulin resistance index (HOMA-IR) = fasting blood glucose (mIU/L) × serum insulin (mmol/L)/22.5.

### Quantitative real-time PCR (RT-qPCR) analysis

(1) RNA isolation: One side of the soleus muscle was taken, and total RNA was extracted with the TRIzol Reagent (Ambion, USA). The RNA concentration and purity were determined with a Nanodrop 2000 nucleic acid analyzer (Thermo Fisher, USA), and the amount of reverse transcription template was calculated based on the RNA concentration. (2) Reverse transcription into cDNA: An All-in-One^TM^ First-Strand cDNA Synthesis Kit (GeneCopoeia, USA) was used with a 25 μL reaction system. The reaction mixture was incubated at 37°C for 1 hour and heated to 85°C for 5 minutes. The reactions were carried out in a Bio-Rad thermal cycle PCR instrument T100. The obtained cDNA product was stored at −20°C. (3) Real-time fluorescence quantification: SYBR^®^ Green All-in-One^TM^ qPCR Mix (GeneCopoeia, USA) was used. The primers used were All-in-One^TM^ qPCR primers supplied by GeneCopoeia, and the uncoupling protein 1 (UCP1) primers were supplied by Bioengineering Co. (Shanghai) (sequences: 5’-gtgtaggcctacaggaccat-3’ and 5’-atgaacatcaccacgttcca-3’). An ABI 7500 realtime PCR instrument was employed. The experimental procedure was as follows: predenaturation at 95°C for 10 minutes, followed by 45 cycles of denaturation at 95°C for 10 seconds, annealing at 60°C for 20 seconds, and extension at 72°C for 15 seconds. All samples were tested 3 times. GAPDH was used as an internal reference, and the results were quantified using the 2^-ΔΔCt^ method.

### Western blotting

A total of 0.1 g of soleus muscle was homogenized on ice, and total protein was extracted with a total protein extraction kit. The protein concentration was measured with the bicinchoninic acid (BCA) method, and 2× loading buffer was used to normalize the concentration according to the measured values. The samples were heated in boiling water for 5 minutes at 100°C, and supernatants were collected for testing after centrifugation. The samples were separated using SDS-PAGE electrophoresis; the concentrations of the stacking and separating gels were determined according to the molecular weights of the target proteins. The proteins were transferred to a polyvinylidene difluoride (PVDF) membrane (Millipore), which was blocked with 5% bovine serum albumin (BSA). The following primary antibodies were used: rabbit anti-hypoxia-inducible factor (HIF)-1α (1:1000, #9475, CST), rabbit anti-peroxisome proliferator-activated receptor gamma coactivator (PGC)-1α (1:1000,ab54481,Abcam), rabbit anti-FGF21 (1:1000, ab171941, Abcam), rabbit anti-UCP1 (1:1000,ab23841, Abcam), rabbit anti-vascular endothelial growth factor A (VEGFA) (1:500, ab9570,Abcam), and rabbit anti-fibronectin type III domain containing 5 (FNDC5) (1:1000, ab131390, Abcam). GAPDH was used as a reference protein, and the primary antibody used was a rabbit anti-GAPDH antibody (horseradish peroxidase (HRP) conjugate, 1:1000,#8884,CST). The membranes were incubated with the primary antibodies overnight at 4°C. The secondary antibody was goat anti-rabbit IgG H&L (HRP) (1:3000, ab205718, Abcam). After incubation with the secondary antibody, the electrochemiluminescence (ECL) solution was applied, and images were collected by a fully automated chemiluminescence image capture system (Tanon 5200). ImageJ 2x software was used to analyze and record the grayscale values of the signal bands.

### Hematoxylin and eosin (HE) staining

Rat inguinal fat embedded in paraffin was sectioned and stained with HE. The specific process was as follows: trimming – fixation – dehydration – transparent – dipping in wax – embedding – slicing – dewaxing – HE staining – sealing. Images were taken using a microscopic imaging system (Olympus, Japan).

### Statistical analysis

SPSS 22.0 and GraphPad Prism 5 were employed for the statistical analysis. Experimental data are expressed as the mean ± standard error. Differences in mean values between two groups were compared using an independent sample T test. When multiple groups of data were homogeneous in variance, one-factor analysis of variance (ANOVA) was used for comparisons among groups. Bonferroni’s multiple comparison test was used for pairwise comparisons. When the variance was not even, Welch’s ANOVA was used, and the Games-Howell test was used for pairwise comparisons (A). The mean values of the rat food intake were compared using variance analysis of repeated measurements. Multifactor variance analysis was used to compare the main effects and interaction effects of hypoxia and endurance exercise. Two-variable correlation analysis employed the Spearman correlation coefficient. p<0.05 indicates a statistically significant difference.

## Results and analysis

### Effects of the HFD on body composition, BG, and blood lipids in rats

Tables 1 and 2 showed that after 8 weeks of the HFD, the average body weight and Lee’s index of the rats in group HFD were higher than those in group N (p<0.05). The average body lengths of the rats were also higher in group HFD than in group N, but the difference was not significant (p>0.05). The BG, TG, TC, and LDL-C levels in the HFD group were higher than those in group N (p<0.05), whereas the HDL-C was significantly lower than that in group N (p<0.05).

**Table 1.**
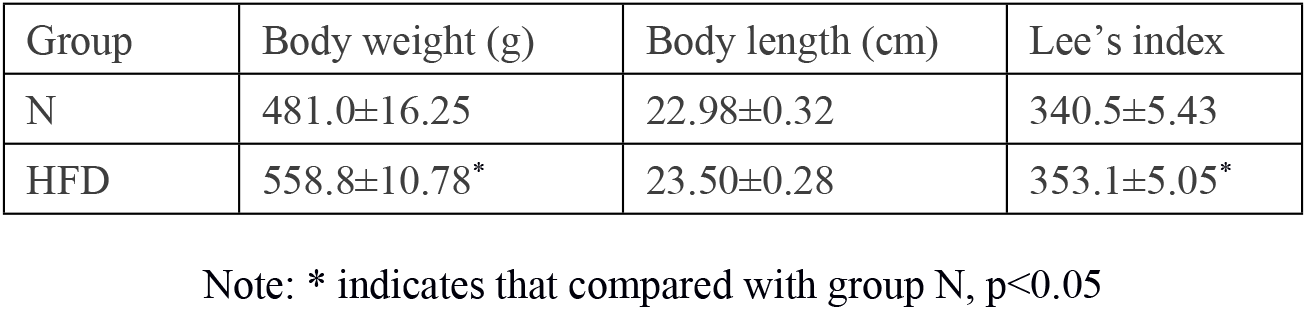
Comparison of body weight and Lee’s index

| Group | Body weight (g) | Body length (cm) | Lee's index |
| --- | --- | --- | --- |
| N | 481.0±16.25 | 22.98±0.32 | 340.5±5.43 |
| HFD | 558.8±10.78* | 23.50±0.28 | 353.1±5.05* |

Note: \* indicates that compared with group N, $p<0.05$

**Table 2.** Comparison of BG and blood lipids

| Group | Blood glucose (BG) | Triglycerides (TGs) | Total cholesterol (TC) | Low-density lipoprotein cholesterol (LDL-C) | High-density lipoprotein cholesterol (HDL-C) |
| --- | --- | --- | --- | --- | --- |
| N | 4.48±0.11 | 1.31±0.12 | 1.25±0.13 | 1.42±0.08 | 0.51±0.05 |
| HFD | 5.75±0.12* | 1.99±0.14* | 2.97±0.14* | 2.08±0.09* | 0.38±0.03* |

Note: \* indicates that compared with group N, $p<0.05$

### Effects of hypoxic training on weight loss in obese rats

As shown in Figure 1A, the body weights of the rats in group A continued to increase during the intervention. The body weights in groups B, C, and D were still increasing in the first 4 weeks. The increase in the rate of body weight decreased from the 5th week, but without a significant difference compared with group A (p>0.05), indicating that hypoxia exposure inhibited the increased rate of body weight gain in the rats fed the HFD to a certain extent, but the effect was not obvious in the short term. The body weight decreased in the first 4 weeks in the AE group but increased from the 5th week. The body weights continued to reduce from the 2nd week in the BE, CE, and DE groups and were significantly different from those in group A (p<0.05). From the 6th week, the body weights of the CE and DE groups were significantly lower than those of the AE group (p<0.05). The results suggest that endurance exercise intervention alone can reduce the body weights of rats fed an HFD in a short period of time but that the body weight rebounds in the later stage. Conversely, hypoxic training continuously reduced the body weights of the obese rats, and the training effects were more significant at the 13.3% and 11.3% oxygen concentrations. Figure 1B shows that the food intake patterns change after 1 week of intervention. The food intake of group A first increased and then decreased, whereas that of group AE first decreased and then increased. The food intakes of groups B, C, and D were lower than that of group A after 5 weeks of intervention, and group C showed a continuous decline. The food intakes of groups BE, CE, and DE were lower than that of group AE at the 8th week, with that of CE group showing a decrease throughout the intervention process. Both hypoxia and hypoxic training could inhibit the food intake of obese rats to varying degrees, with 13.3% hypoxic training having the largest effect. Exercise alone could reduce food intake in the early stage, but food intake increased again in the later stage.

**Figure 1.**
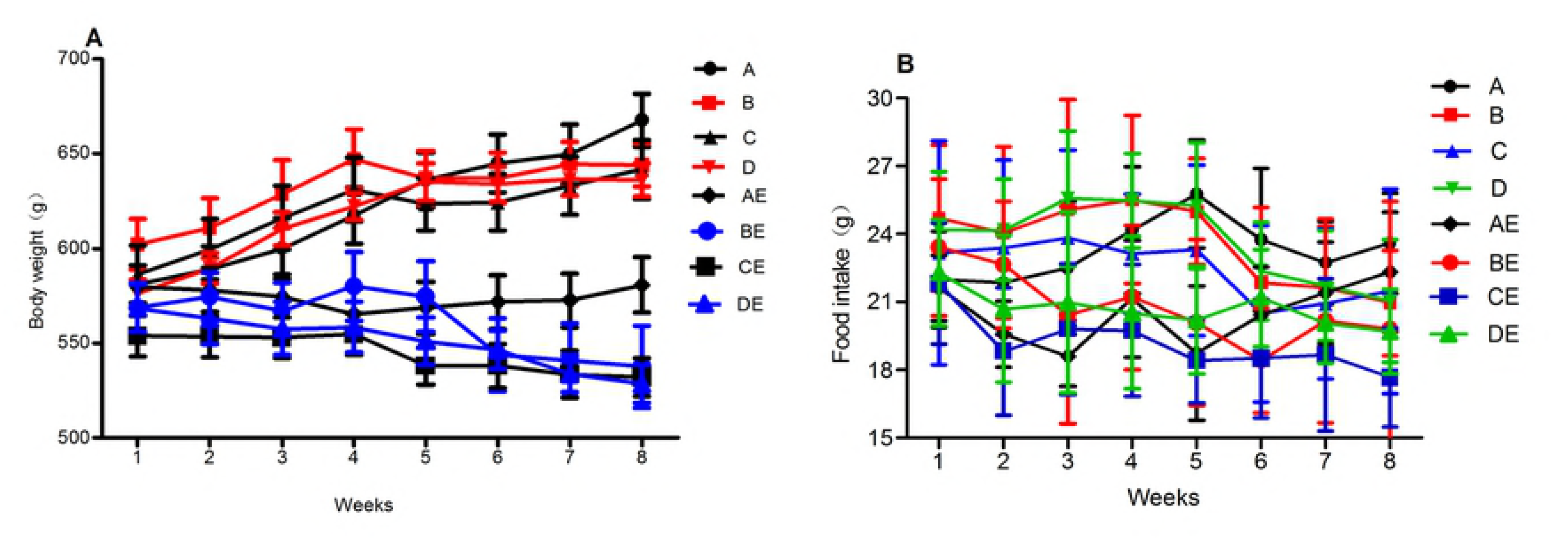
Changes in body weight and food intake during the hypoxic training intervention

**Figure 2.**
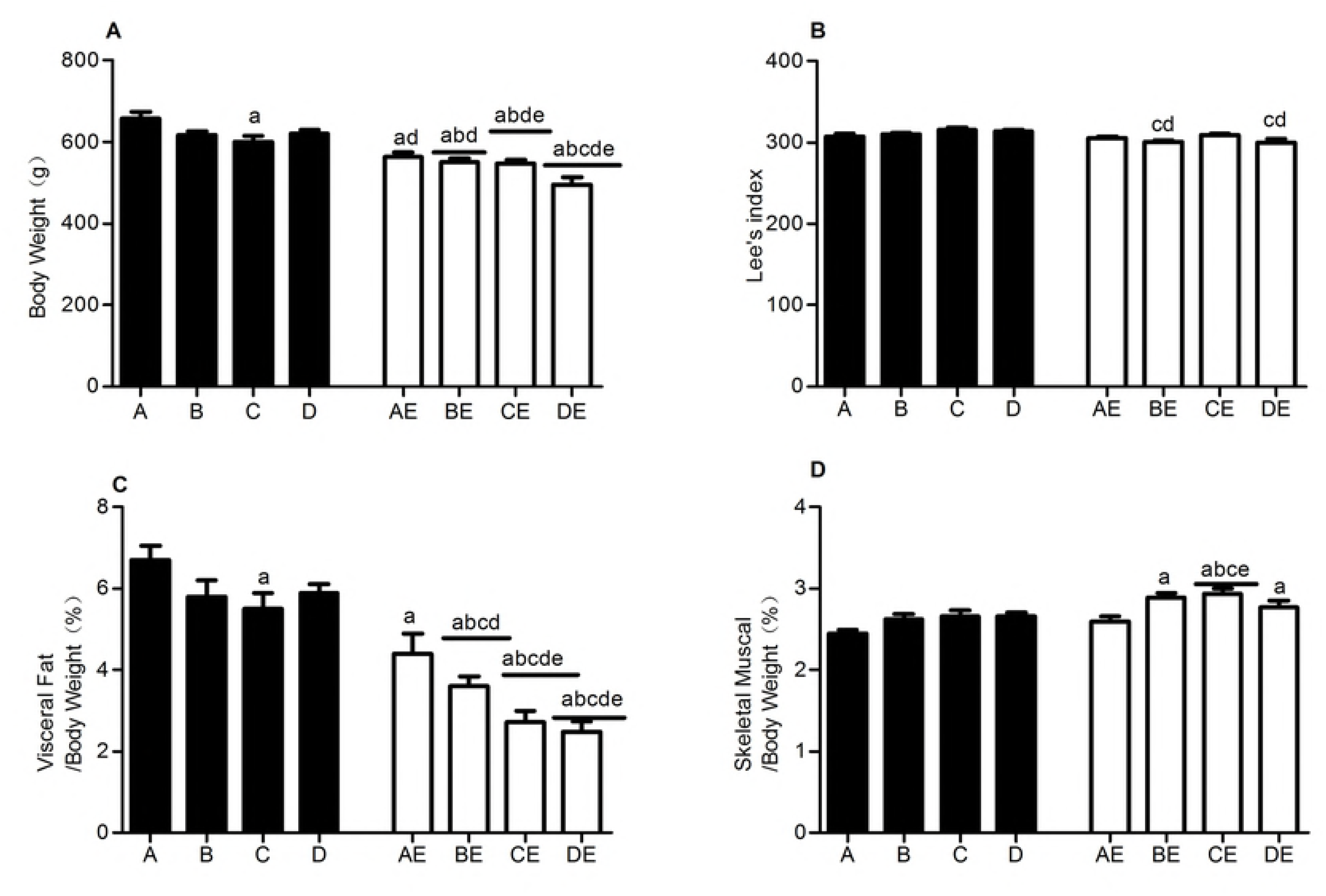
HE staining of inguinal adipose tissue after the interventions

After 8 weeks of intervention, the group A rats had the fewest fat cells in the same microscopic field of inguinal adipose HE-stained tissue compared with those of the other groups and the largest area of fat droplets in single fat cells. Compared with the AE group, the BE, CE, and DE groups had more fat cells and smaller lipid droplets, with the most significant difference observed in the DE group. The BE, CE, and DE groups had more fat cells and smaller lipid droplets than the B, C and D groups, respectively. The results suggest that hypoxia exposure, endurance exercise, and hypoxic training can improve the morphology of fat cells and reduce lipid droplets. The lower oxygen concentrations resulted in more obvious intervention effects, and hypoxic training was more effective than hypoxia exposure or endurance training alone.

Note: a represents compared with group A, p<0.05; b represents compared with group B, p<0.05; c represents compared with group C, p<0.05; d represents compared with group D, p<0.05; and e represents compared with group AE, p<0.05.

After 8 weeks of intervention, the body weights of group A were the highest. The body weights of groups C, AE, BE, CE, and DE were significantly lower than those of group A (p<0.05). No significant difference in body weight was observed among groups B, C, and D (p>0.05), but all of the body weights in these groups were higher than those of group AE (p<0.05). The body weights of groups BE, CE, and DE were lower than those of group AE, and those of groups CE and DE were significantly different from those of group AE (p<0.05). The DE group had the lowest body weights (Fig 3A). These results indicate that 13.3% hypoxia exposure can significantly reduce the body weights of obese rats fed an HFD, exercise alone is better than hypoxia exposure, the effect of hypoxic training is better than that of hypoxia exposure or endurance training, and a lower oxygen concentration has a more significant effect. Lee’s indexes of the rats in groups C and D were significantly higher than those in groups BE and DE (p<0.05), suggesting that hypoxia exposure or endurance exercise did not effectively reduce Lee’s index in obese rats, whereas 16.3% and 11.3% hypoxic training was effective in reducing Lee’s index in obese rats (Fig 3B). The percentages of visceral fat in the rats in groups C, AE, BE, CE, and DE were significantly lower than that in group A (p<0.05); the percentage was lowest in group DE, and the percentages in groups BE, CE, and DE were significantly lower than that in group AE (p<0.05) and those in the corresponding hypoxia exposure groups (p<0.05). The above results indicate that hypoxia and/or exercise can reduce the percentage of visceral fat to body weight. The effect of 13.3% hypoxia exposure was better than that of 16.3% and 11.3% hypoxia exposure. Hypoxia combined with endurance exercise was more effective than hypoxia exposure or exercise alone, and the lower oxygen concentrations had more significant effects (Fig 3C). The percentage of skeletal muscle to body weight of group A was lowest, and that of group CE was highest. Those of groups BE, CE, and DE were significantly increased compared with that of group A (p<0.05). The percentage of skeletal muscle to body weight in group CE was significantly higher than that in group AE (p<0.05), indicating that both hypoxia exposure and exercise could increase the percentage of skeletal muscle to body weight, with 13.3% hypoxic training showing the best effect (Fig 3D). A comprehensive review of the body weight, visceral fat percentage, and skeletal muscle percentage indicated that the decrease in body weight was mainly due to the decrease in the visceral fat content.

**Figure 3.**
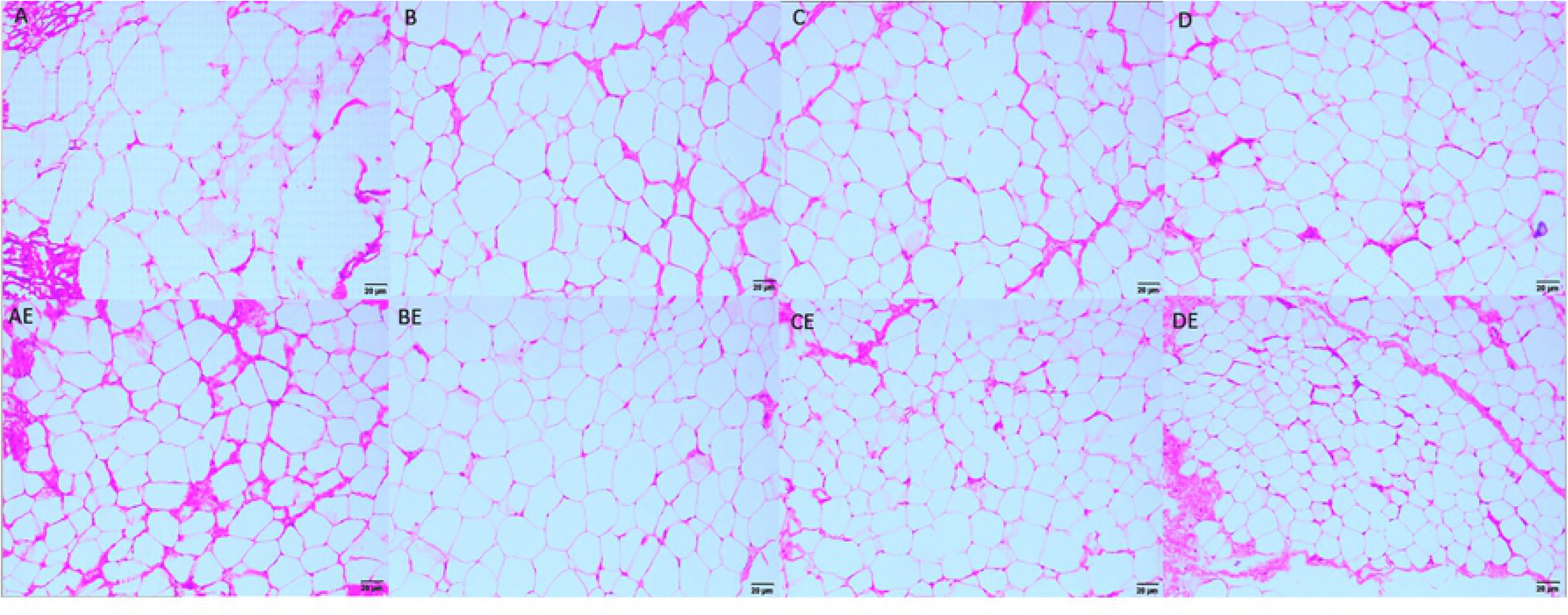
Comparison of the rat body compositions after the intervention

**Figure 4.**
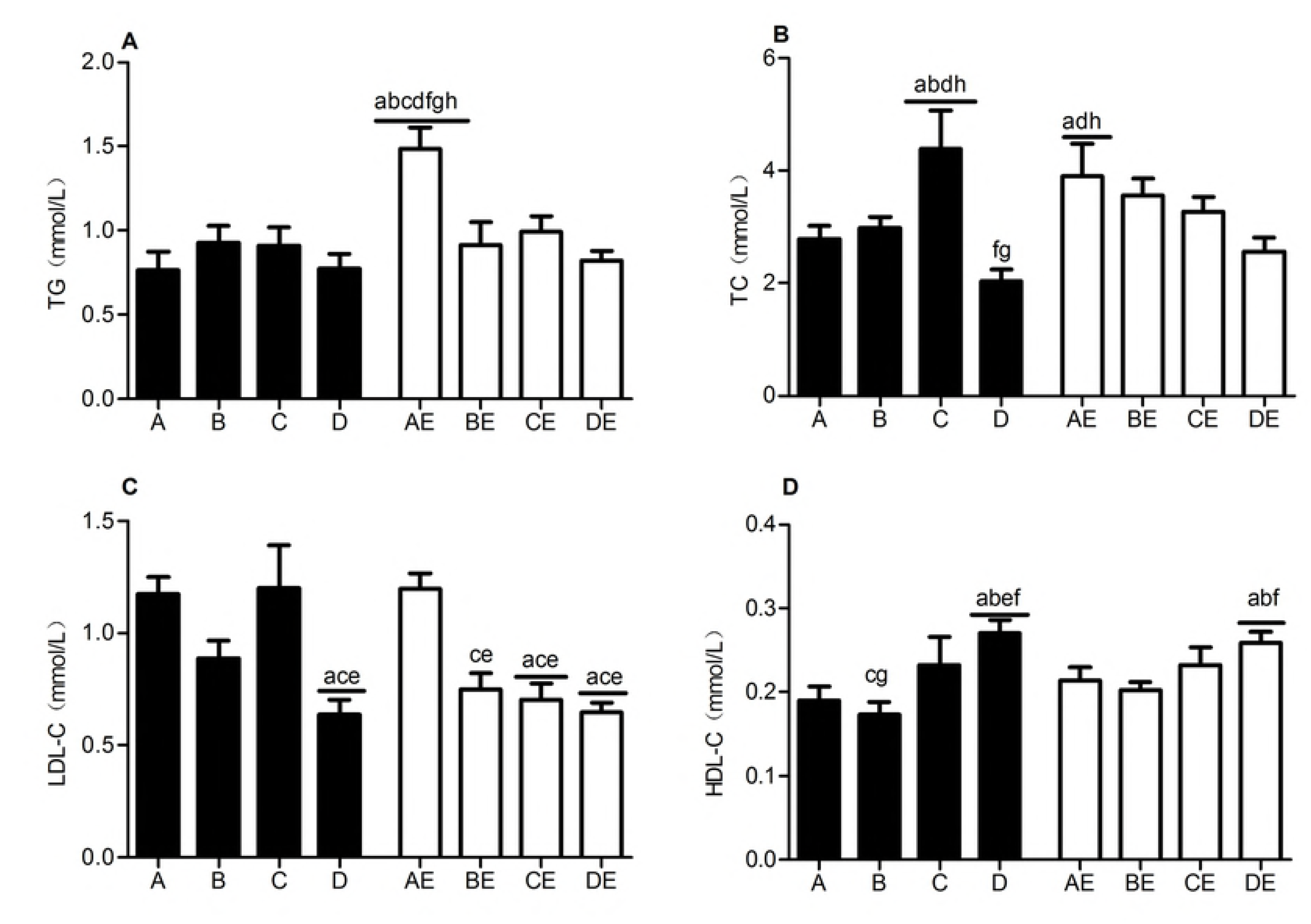
Comparison of blood lipids after the intervention

Note: a represents compared with group A, p<0.05; b represents compared with group B, p<0.05; c represents compared with group C, p<0.05; d represents compared with group D, p<0.05; e represents compared with group AE, p<0.05; and f represents compared with group BE, p<0.05.

The AE group had the highest TG level (p<0.05). The serum total cholesterol (TC) content was highest in group C and lowest in group D, whereas those in the AE, BE, CE, and DE groups showed a decreasing trend. The LDL-C contents were significantly higher in groups D, BE, CE, and DE than in groups A and AE and showed a decreasing trend in the order of groups AE, BE, CE, and DE. The HDL-C levels in groups D and DE were significantly higher than that in group A, and the level in group D was also higher than that in group AE (p<0.05). The above results indicated that after 8 weeks of intervention, endurance exercise significantly increased the serum TG levels in the obese rats, 13.3% hypoxia exposure caused a significant increase in CHO, and 11.3% hypoxia exposure and hypoxic training significantly reduced the TG, CHO, and LDL-C contents and elevated the HDL-C concentration.

### Effects of hypoxic training on the serum irisin and adiponectin levels in obese rats

Note: a represents compared with group A, p<0.05; b represents compared with group B, p<0.05; c represents compared with group C, p<0.05; d represents compared with group D, p<0.05; e represents compared with group AE, p<0.05; and g represents compared with group CE, p<0.05.

The serum irisin content was higher in group CE than in groups A, C, and D (p<0.05). The serum irisin content of group DE was significantly higher than those of groups A, B, C, D, and AE (p<0.05). Hypoxia exposure or endurance training alone did not increase the serum irisin levels, whereas 13.3% and 11.3% oxygen concentration training showed significant effects (Fig 5A). The serum adiponectin concentrations of groups C, D, AE, BE, and DE were lower than those of groups A and B (p<0.05). The 13.3% and 11.3% hypoxia exposures reduced the adiponectin concentration, and endurance training and 16.3% and 11.3% hypoxic training also reduced the adiponectin concentration in the rat sera (Fig 5B). The leptin level was higher in group A than in the other groups (p<0.05), was lower in group D than in groups B and C (p<0.05), and was lower in group DE than in groups AE and BE (p<0.05). Hypoxia exposure and/or exercise reduced the serum leptin levels, with 11.3% hypoxia exposure and hypoxic training showing the most pronounced effects (Fig 5C). The HOMA-IR was significantly lower in groups B, C, D, BE, CE, and DE than in group A (p<0.05) and was lower in groups CE and DE than in group AE (p<0.05), indicating that both hypoxia exposure and hypoxic training at the three oxygen concentrations could improve IR in obese rats. Additionally, training at the 13.3% and 11.3% oxygen concentrations improved IR in obese rats better than endurance training alone (Fig 5D).

**Figure 5.**
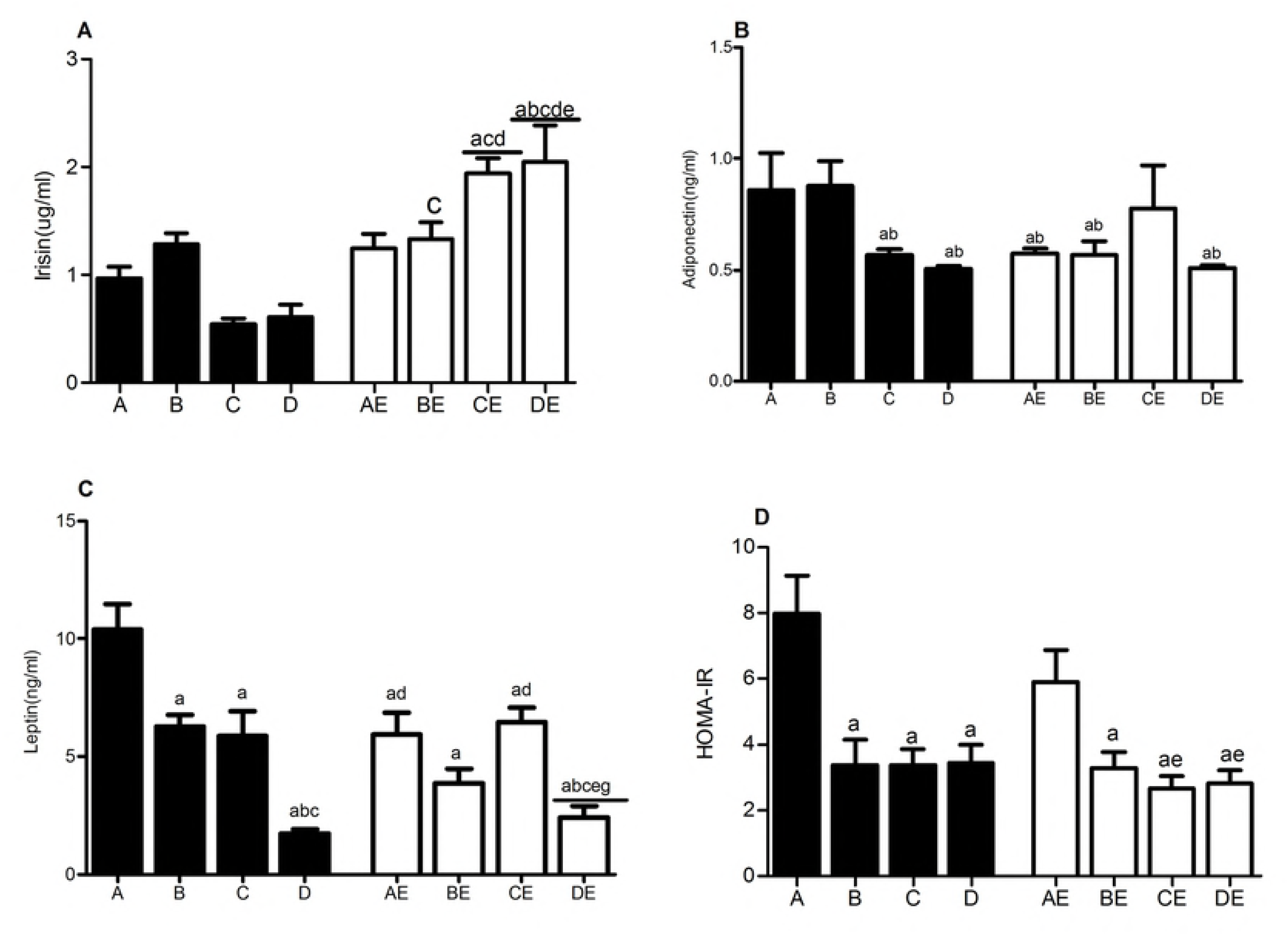
Comparison of the rat serum irisin, adiponectin, and leptin levels and the HOMA-IR

### Effects of hypoxic training on the transcription and protein expression levels of index genes related to browning of white fat

Relative HIF-1α mRNA expression in the skeletal muscle was significantly lower in groups A and AE than in the other 6 groups (p<0.05). Hypoxia exposure and hypoxic training promoted HIF-1α mRNA expression in the skeletal muscle. The HIF-1 α mRNA transcription level was inversely proportional to the oxygen concentration (Fig 6A). The HIF-1α mRNA and protein contents were significantly higher in groups D and DE than in group A (p<0.05), with 11.3% hypoxia exposure and/or endurance training showing the most obvious effects (Fig 6A and 7A).

**Figure 6.**
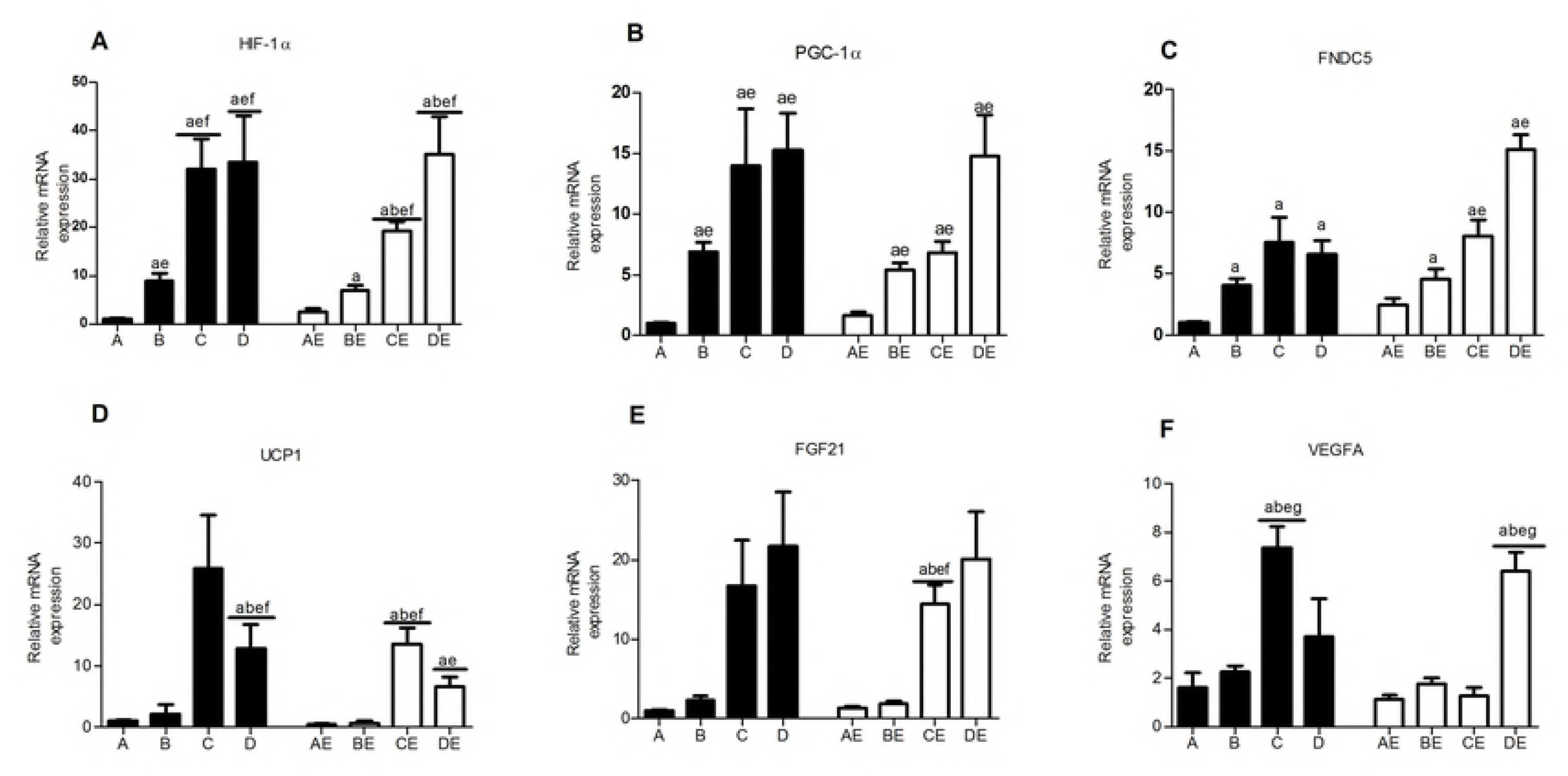
Expression of browning-related genes after the intervention

**Figure 7.**
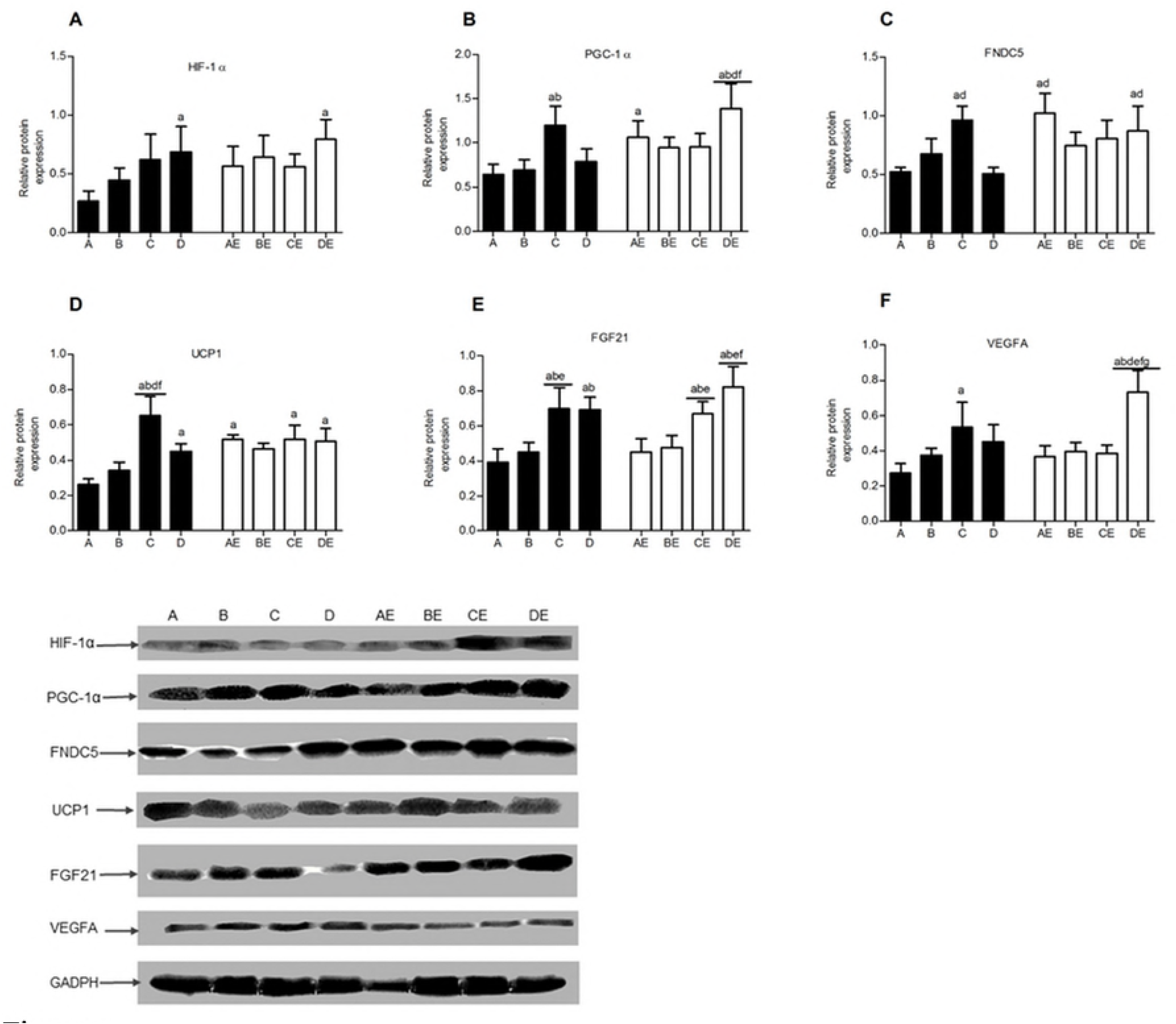
Expression of browning-related proteins after the intervention

The three different concentrations of hypoxia exposure and hypoxic training significantly increased PGC-1α and FNDC5 gene expression in the skeletal muscle. The lower oxygen concentrations resulted in higher mRNA transcription levels. Conversely, the effect of endurance exercise alone was not obvious. The DE group showed the highest FNDC5 gene expression level (Fig 6B and 6C). The PGC-1α protein contents in the skeletal muscle were significantly higher in groups C, AE, and DE than in group A (p<0.05). The level in group DE was the highest and was significantly different from that in groups A, B, D, and BE (p<0.05) (Fig. 7B). FNDC5 protein expression was higher in groups C, AE, and DE than in groups A and D (p<0.05), indicating that 13.3% hypoxia exposure, endurance exercise alone, and 11.3% hypoxic training could promote FNDC5 protein expression in the skeletal muscle of obese rats (p<0.05) (Fig 7C). The UCP1 mRNA transcription level was significantly higher in groups C, D, CE, and DE than in groups A and AE (p<0.05), indicating that 13.3% and 16.3% hypoxia exposure and hypoxic training promoted UCP1 gene transcription, whereas exercise alone had no obvious effect on UCP1 gene expression (Fig 6D). The UCP1 protein expression level in the skeletal muscle in group A was lower than that in the other 7 groups and demonstrated significant differences compared with those in groups C, D, AE, CE, and DE (p<0.05), indicating that both hypoxia exposure and hypoxic training could increase UCP1 protein expression, although the 13.3% and 11.3% hypoxic environments and hypoxic training were more effective (Fig 7D). The FGF21 gene transcription and protein expression levels in groups C, D, CE, and DE were significantly higher than those in groups A and AE (p<0.05) (Fig 6E and 7E), indicating that 13.3% and 11.3% hypoxia exposure and hypoxic training significantly increased FGF21 gene transcription and protein expression in skeletal muscle. VEGFA gene transcription and protein expression were significantly higher in groups C and DE than in group A and were significantly higher in group DE than in groups AE, BE, and CE (p<0.05) (Fig 6F and 7F), indicating that the VEGFA gene and protein expression levels were highly consistent, 13.3% hypoxia exposure and 11.3% hypoxic training showed significant effects on VEGFA expression, and 11.3% hypoxic training was superior to endurance training at the other low oxygen concentrations.

### Correlation analysis of the serum irisin concentration in the rats

Table 3 shows that the serum irisin concentration after 8 weeks of intervention was significantly negatively correlated with the body weight, Lee’s index, and the visceral fat percentage (p<0.05) and significantly positively correlated with the skeletal muscle percentage and FNDC5 mRNA transcription level (p<0.05).

**Table 3.**
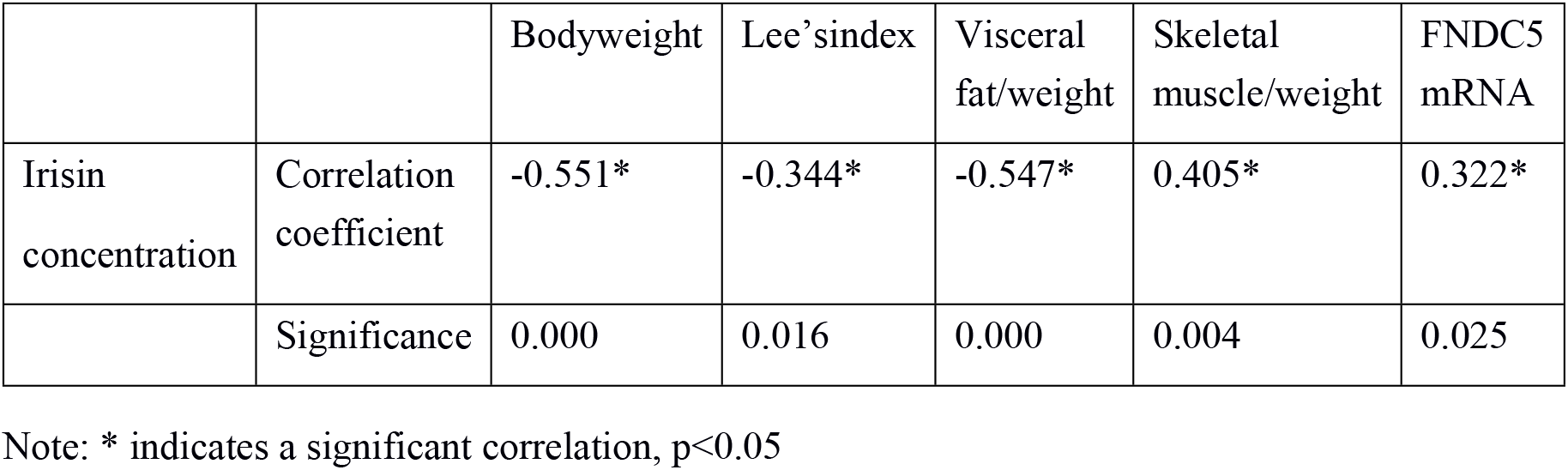
Two-factor correlation analysis

|  |  | Bodyweight | Lee's index | Visceral fat/weight | Skeletal muscle/weight | FNDC5 mRNA |
| --- | --- | --- | --- | --- | --- | --- |
| Irisin concentration | Correlation coefficient | -0.551* | -0.344* | -0.547* | 0.405* | 0.322* |
|  | Significance | 0.000 | 0.016 | 0.000 | 0.004 | 0.025 |

Note: \* indicates a significant correlation, $p<0.05$

## Discussion

### Effects of hypoxia exposure on weight loss in obese rats

HFD feeding causes excessive fatty acids in rats, which inhibit the secretion of various regulatory hormones and utilization of glucose by skeletal muscle. Local hypoxic necrosis of tissue caused by excessive fat accumulation is an important cause of insulin resistance, leading to abnormal glucose and lipid metabolism in obese rats[25, 26]. The oxygen supply plays an important role in regulating body weight and energy metabolism[27]. Human and animal experiments have confirmed that hypoxic environments can alter body composition and can be used as an intervention and treatment for obesity and related metabolic diseases[28] However, hypoxia exposure is a double-edged sword, especially under the condition of a relatively low oxygen concentration when the loss of protein is significant and the damage to the body cannot be ignored[29]. Therefore, the ideal state of hypoxic intervention in weight loss improves utilization of fat and avoids loss of skeletal muscle[30].

Our study found that hypoxia exposure alone inhibited the excessive growth of body weight to some extent, but the body weight still showed an upward trend compared with the weight before the intervention. Of the three hypoxia exposure concentrations, 13.3% showed the most significant effects. At the end of the 8-week intervention, the body weights of the 13.3% hypoxia exposure group were significantly lower than those of the quiet control group. Therefore, among the three low oxygen concentrations, the moderate 13.3% oxygen concentration demonstrated the best control effect on body weight. In addition, the visceral fat percentage of the rats in the 13.3% hypoxia exposure group was lower than those in the other two groups and was significantly lower than that in the quiet control group. Moreover, the percentage of skeletal muscle in this group was higher than those in the other three groups without exercise intervention, although the difference was not significant. Lee’s index showed no obvious difference, indicating that 13.3% hypoxia exposure mainly increased the energy supply by fat and did not significantly decompose proteins. Previous studies suggested that during weight loss caused by high altitude hypoxia exposure, protein synthesis of skeletal muscle was inhibited, decomposition was enhanced, and protein loss was significan[31–34]. However, under these situations, the oxygen concentrations of hypoxia exposure were low (altitude >5000 m), and the hypoxia mode was continuous exposure. The results of this study indicate that intermittent hypoxia exposure at a suitable concentration does not cause loss of skeletal muscle proteins. The three hypoxia exposure concentrations showed significant effects on food intake at later stages of the intervention, and the energy intake in the latter four weeks was lower than that of the quiet control group. Therefore, hypoxia exposure alone demonstrated a certain weight loss effect on obese SD rats. One possible reason may be that hypoxia exposure resulting in tissue anoxia inhibits the appetites of rats, increases fat hydrolysis and mobilization, and increases the energy output, which are similar to the conclusions of previous studies[17, 35]. However, after 8 weeks of intervention, the serum leptin concentration was significantly lower in the hypoxia exposure groups than in the quiet control group, which was inconsistent with the results reported in some studies [36, 37]. A possible explanation is that the obesity caused by the early HFD leads to leptin resistance in the rats, and hypoxia exposure improves this condition. The changes in the HOMA-IR in each group after 8 weeks also suggest that intermittent hypoxia exposure improves the condition of IR, which is similar to the hypoxia exposure model established by Chen et al[16] with hypoxia exposure for 8 hours per day and six days per week to an oxygen concentration of 14-15%. Moreover, the serum TG and CHO levels in the hypoxia exposure groups were not significantly different from those in the quiet group, whereas the serum LDL-C concentration was significantly decreased and the HDL-C concentration was significantly increased in the 11.3% oxygen concentration group. Hypoxia exposure showed a more obvious weight loss effect for individuals with higher body weights[38], with lower oxygen concentrations leading to more weight loss. However, obese individuals are more prone to mountain sickness; therefore, when low-oxygen exposure is applied for the purpose of weight loss, attention should be paid to the mode of hypoxia exposure and the choice of oxygen concentration. In conclusion, 13.3% intermittent hypoxia exposure has a significant effect on weight loss in nutritionally obese rats, and 11.3% hypoxia interventions can improve the state of metabolic disorders.

### Effects of hypoxic training on weight loss in obese rats

Hypoxic training can realize regulation of the body’s metabolic capacity through low oxygen exposure and exercise[39], avoid the risk of hypoxia exposure through appropriate exercise training[40], and achieve weight loss optimization [41]. In previous reports, the choice of an oxygen concentration for safe and effective weight loss was mainly concentrated at a simulated altitude of 1700 m-5000 m (oxygen concentration of 11.3%-17.3%), and hypoxia exposure was mainly based on an intermittent hypoxia intervention combined with endurance exercise and moderate resistance training[41]. During the 8-week intervention, the inhibition of weight gain by endurance training alone was better than that of hypoxia exposure, whereas the effect of hypoxic training was better than that of hypoxia exposure or endurance exercise, and the intervention became more effective as the oxygen concentrations decreased. Hypoxic training demonstrated a significant inhibitory effect on food intake. Endurance exercise alone inhibited food intake in the early stage, but food intake showed a continuously increasing trend in the later stage. This phenomenon may occur because the animals are not adapted to the exercise load at the early stage, and thus, exercise fatigue inhibits their food intake; after the animals are adapted in the later stage, the food intake gradually increases, suggesting that exercise has little effect on appetite[42]. However, hypoxic training could continuously inhibit food intake, and the food intake amount demonstrated a trend of a continuous decrease. Endurance training with different oxygen concentrations significantly reduced the body weights and visceral fat percentages in obese rats and increased the percentages of skeletal muscle mass. The lower oxygen concentrations had more obvious effects. The 11.3% hypoxic training demonstrated the most significant effects on weight control. The 16.3% and 13.3% hypoxic training not only increased the energy supply by fat decomposition but also promoted protein synthesis. However, training at the 11.3% oxygen concentration resulted in a lower percentage of skeletal muscle mass, suggesting that the energy supply from the protein proportion might be higher than that in the other two groups. Both endurance and hypoxic training increased the serum TG and CHO concentrations, probably because under continuous feeding of the HFD, endurance training and/or hypoxic training enhanced the usage of stored fat and mobilized fat into the bloodstream, thereby improving the efficiency of fat recycling. Both 13.3% and 11.3% hypoxic training increased the serum HDL concentration and decreased the LDL level, which was a sign of improvement in lipid metabolism. Hypoxic training reduced serum leptin and HOMA-IR to varying degrees and improved leptin and IR in obese rats, similar to the results of previously reported animal intervention studies[43–45]. Hypoxic training can effectively act on skeletal muscle and oxygen responses, resulting in adaptation of health-related molecules at the skeletal muscle level. Longer hypoxic training has even more obvious effects. Therefore, long-term hypoxic training may be used as an effective nonpharmacological method to treat and prevent IR.

### Effects of hypoxic training on the circulating irisin concentration

Irisin is a myokine[2] and adipokine[46]that is released after PGC-1α activation and is edited from FNDC5. Irisin has received wide attention because it promotes white fat browning, improves the body’s energy output without changing the exercise and diet status, and improves the obesity condition and glycolipid metabolism. Studies have found that irisin upregulates UCP1 expression, possibly through p38MAPK and ERK phosphorylation[47].

Currently, it is generally agreed that irisin promotes energy metabolism and improves obesity and IR in rodents[48]. However, study results concerning whether the irisin concentration increases in human patients with metabolic diseases are inconsistent. Some studies found that the irisin level in the blood of patients with type 2 diabetes (T2D) was significantly lower than that of people with normal BG[49, 50]. FNDC5 gene expression was significantly decreased in skeletal muscle of obese people, and FNDC5 expression was significantly decreased in subcutaneous and visceral fat[51]. Other studies found that the baseline levels of blood irisin in patients with obesity, metabolic syndromes, and T2D were significantly higher than those in the general population. Irisin was negatively correlated with the adiponectin level and positively correlated with the body mass index, fasting BG, TGs, and IR. The theory of irisin resistance was proposed, which stated that both skeletal muscle and adipose tissue increased irisin secretion to overcome irisin resistance[5, 52, 53]. FNDC5 is downregulated in T2D patients and upregulated in *in vitro* skeletal muscle cell culture, indicating that diabetes-associated factors regulate FNDC5/irisin expression *in vivo*^[53]^. Therefore, the irisin content is related to the skeletal muscle volume and plays a role in glycolipid metabolism. In addition, irisin secretion is mainly the result of the interaction of skeletal muscle and fat tissue. Because FNDC5 expression in skeletal muscle far exceeds that in adipose tissue, 70% of irisin has been postulated to be secreted by skeletal muscles. The secretion of various proteins by skeletal muscle is mainly dependent on its contraction function[54], and thus, physical activities are considered the main factor affecting irisin secretion. However, results from current research regarding whether exercise can promote irisin secretion and whether irisin plays a regulatory role in exercise to improve metabolism of the body are controversial[53]. Meta et al. found that the irisin concentration in humans and animals after various acute training sessions increased under appropriate strength conditions[55, 56]. In addition, experiments found that long-term training promoted irisin expression and an increased concentration in animals[57]. However, this effect has not been observed in human experiments[55, 56]. Within 1-2 hours after exercise, the irisin concentration in obese people reached the highest peak[58]. Changes in the irisin concentration were mainly related to the ATP levels, followed by glucose and lipid metabolism in skeletal muscle[59, 60].

Our results showed that hypoxia exposure interventions demonstrated certain inhibitory effects on the serum irisin concentration of rats, but no significant difference was found compared with the quiet control group, which was similar to the results of previous human studies[61, 62]. Endurance training alone had little effect on the rat irisin concentration, and no significant difference was found compared with the quiet control group. Long-term endurance training did not increase the irisin concentration in obese rats, possibly because the samples were not measured immediately after the last training session[58]. After hypoxic training, the irisin concentration increased. Multifactor variance analysis confirmed that hypoxia had no significant effect on the irisin concentration, whereas exercise and hypoxic training showed significant effects, with lower oxygen concentrations showing the most obvious effects. The 11.3% hypoxic training group showed effects that were significantly higher than those of the nonexercise and endurance training groups. Correlation analysis found that the serum irisin concentration was significantly correlated with the skeletal muscle mass percentage. The increase in the irisin concentration may be due to the increase in the skeletal muscle mass percentage in rats after hypoxic training. Previous studies identified a negative correlation between the irisin and adiponectin concentrations in blood[5, 49]. This relationship was not observed in this study. Previous studies found that acute anoxia and persistent hypoxia could increase leptin secretion by human white adipose precursor cells but had limited effects on adiponectin[63]. Leptin increases adiponectin expression under normal oxygen conditions, but this effect is not detected under short-term hypoxic conditions. In this study, 13.3% and 11.3% hypoxia exposure reduced the serum adiponectin concentration, which demonstrated a significant positive correlation with the serum leptin concentration, indicating that leptin could affect the adiponectin level under long-term intermittent hypoxia conditions. This finding was similar to the results obtained by Fu C[64]in experiments simulating chronic intermittent hypoxia. The three hypoxia concentrations reduced the leptin concentration and HOMA-IR in obese rats, indicating that hypoxia exposure improved glucose metabolism, increased insulin sensitivity, and decreased leptin resistance in this animal model.

### Effects of hypoxic training on white fat browning in obese rats

HIF-1α is mainly regulated by the oxygen concentration. This protein plays an important role in maintaining heat production by brown fat and promoting mitochondrial proliferation and vascular synthesis[65]. The three hypoxia exposure concentrations used in this study promoted HIF-1α overexpression, with the lower oxygen concentrations promoting higher HIF-1α mRNA transcription levels. Furthermore, hypoxic training did not aggravate the hypoxia state of the body compared with hypoxia exposure. HIF-1α protein expression was significantly affected by hypoxia exposure and was significantly higher in the 11.3% hypoxia exposure group and 11.3% hypoxic training group than in the quiet control group. In addition, the PGC-1α, FNDC5, UCP1, FGF21, and VEGFA mRNA transcription levels were affected by hypoxia exposure. The PGC-1α mRNA expression pattern was similar to that of HIF-1α, indicating that HIF-1α might be one upstream regulator of PGC-1α. The FNDC5 mRNA transcription level was not significantly different among the three hypoxia concentrations. Hypoxia exposure may ultimately affect FNDC5 transcription as an indirect influencing factor. The mRNA transcription levels of genes related to white fat browning (UCP1, FGF21, and VEGFA) were significantly affected by the low oxygen concentrations, suggesting that low oxygen concentrations during hypoxia exposure might promote the expression of brown fat transformation-related genes in obese rats.

Endurance training improved PGC-1α and FNDC5 gene expression, but this expression showed no significant difference compared with that in the quiet control group, indicating that long-term endurance training had no obvious effect on the expression of white fat browning-related genes, possibly because the body adapted to long-term endurance training. However, PGC-1α, FNDC5, and UCP1 protein expression in the skeletal muscle of obese rats was significantly increased after endurance training. Factorial analysis found that exercise was the main cause of regulated expression of the three proteins, and thus, other pathways might improve their translation levels. These results are consistent with those of De Matteis, R et al. [66], indicating that endurance training can be used as a method to stimulate browning of white fat. The HIF-1α and PGC-1α transcription levels were significantly higher in the hypoxic training groups than in the quiet control group and were negatively correlated with the oxygen concentration. The oxygen concentration was identified as the main effector, indicating that these two genes were mainly stimulated by hypoxia. The FNDC5 mRNA level was higher in all hypoxic training groups than in the quiet control group and showed a different change trend in the hypoxic training groups than under hypoxia exposure alone, indicating that hypoxic training achieved the dual effects of hypoxia exposure and exercise training. The PGC-1α and FNDC5 protein expression levels were significantly higher in the endurance training groups and the 11.3% hypoxic training group than in the quiet control group, indicating that endurance training was more effective than hypoxia exposure and that hypoxia exposure to lower oxygen concentrations combined with exercise was more effective. UCP1 expression was higher in the 4 exercise groups than in the quiet control group. The effect of hypoxic training was not significantly different from that in the exercise group alone, indicating that endurance training was the main factor that promoted UCP1 expression. Both 13.3% and 11.3% hypoxic training significantly increased the FGF21 protein expression level. As a motility factor, FGF21 can regulate energy metabolism through the AMPK-SIRTl-PGClα pathway[67], mainly by increasing cell lipid oxidation to provide energy, promoting glucose uptake by the liver and kidney, enhancing insulin resistance, and promoting white fat browning[68]. VEGFA has functions of maintaining vascular density in brown adipose tissue and improving the IR of obese individuals[69]. Training under the 11.3% oxygen concentration demonstrated the most significant effect on promoting VEGFA protein expression, indicating that this hypoxic training concentration was a suitable trigger condition. In summary, hypoxia training showed a more significant effect in promoting transcription and translation of white fat browning-related genes than hypoxia exposure or endurance training alone, with training under the 11.3% oxygen concentration demonstrating the most significant effect.

## Conclusions

Hypoxia exposure, endurance training, and hypoxic training demonstrated effects of promoting weight loss and improving glucose and lipid metabolism. The effects of hypoxic training were more significant than those of hypoxia exposure or endurance training alone, with 13.3% hypoxia exposure and 11.3% hypoxic training having the best effects.

Hypoxia exposure or endurance training alone did not significantly change the serum irisin concentration. Hypoxic training increased the irisin concentration in nutritionally obese rats, and hypoxic training at the 11.3% oxygen concentration showed the most obvious effects.

Hypoxia exposure and hypoxic training showed various degrees of promotion of transcription of genes related to white fat browning. Endurance training and hypoxic training promoted protein expression, with 11.3% hypoxic training showing the most significant effect.

## Supporting Information

S1 Table. The results of factorial analysis

## References

1. Abarca-Gómez L, Abdeen ZA, Hamid ZA, Dika Z, Ivkovic V, Jelakovic A, et al. Worldwide trends in body-mass index, underweight, overweight, and obesity from 1975 to 2016: a pooled analysis of 2416 population-based measurement studies in 128·9 million children, adolescents, and adults. Lancet. 2017;390(10113):2627–42. doi:10.1016/S0140-6736(17)32129-3.

2. Bostrom P, Wu J, Jedrychowski MP, Korde A, Ye L, Lo JC, et al. A PGC1-alpha-dependent myokine that drives brown-fat-like development of white fat and thermogenesis.Nature.2012;481(7382):463–U72. doi:10.1038/nature10777 PMID: 22237023; PubMed Central PMCID:PMC3522098.

3. Polyzos SA, Mathew H, Mantzoros CS. Irisin: A true, circulating hormone. Metabolism. 2015;64(12):1611–8. doi:10.1016/i.metabol.2015.09.001 PMID: 26422316.

4. Hecksteden A, Wegmann M, Steffen A, Kraushaar J, Morsch A, Ruppenthal S, et al. Irisin and exercise training in humans – Results from a randomized controlled training trial. BMC Med. 2013;11:8. doi: 10.1186/1741-7015-11-235 PMID: 24191966.

5. Park KH, Zaichenko L, Brinkoetter M, Thakkar B, Sahin-Efe A, Joung KE, et al. Circulating Irisin in Relation to Insulin Resistance and the Metabolic Syndrome. J Clin Endocrinol Metab. 2013;98(12):4899–907. doi:10.1210/jc.2013-2373 PMID: 24057291.

6. Arias-Loste MT, Ranchal I, Romero-Gomez M, Crespo J. Irisin, a Link among Fatty Liver Disease, Physical Inactivity and Insulin Resistance. Int J Mol Sci. 2014;15(12):23163–78. doi:10.3390/ijms151223163 PMID: 25514415.

7. Huh JY, Siopi A, Mougios V, Park KH, Mantzoros CS. Irisin in Response to Exercise in Humans With and Without Metabolic Syndrome. J Clin Endocrinol Metab. 2015;100(3):E453–E7. doi:10.1210/jc.2014-2416 PMID: 25514098.

8. Vamvini MT, Hamnvik OP, Sahin-Efe A, Gavrieli A, Dincer F, Farr OM, et al. Differential Effects of Oral and Intravenous Lipid Administration on Key Molecules Related to Energy Homeostasis. J Clin Endocrinol Metab. 2016;101(5):1989–97. doi:10.1210/jc.2015-4141 PMID: 26964729.

9. Huh JY, Mantzoros CS. Irisin physiology, oxidative stress, and thyroid dysfunction: What next? Metabolism. 2015;64(7):765–7. doi:10.1016/j.metabol.2015.02.009 PMID: 25916681.

10. Assyov Y, Gateva A, Tsakova A, Kamenov Z. Irisin in the Glucose Continuum. Exp Clin Endocrinol Diabet. 2016;124(1):22–7. doi:10.1055/s-0035-1564130 PMID: 26479549.

11. Qiu SH, Cai X, Sun ZL, Schumann U, Zugel M, Steinacker JM. Chronic Exercise Training and Circulating Irisin in Adults: A Meta-Analysis. Sports Med. 2015;45(11):1577–88. doi:10.1007/s40279-014-0293-4 PMID: 26392122.

12. Bell MA, Levine CB, Downey RL, Griffitts C, Mann S, Frye CW, et al. Influence of endurance and sprinting exercise on plasma adiponectin, leptin and irisin concentrations in racing Greyhounds and sled dogs Aust Vet J. 2016;94(5):154–9. doi:10.1111/avj.12436 PMID: 27113986.

13. Chen N, Li QX, Liu J, Jia SH. Irisin, an exercise-induced myokine as a metabolic regulator: an updated narrative review Diabetes-Metab Res Rev. 2016;32(1):51–9. doi:10.1002/dmrr.2660 PMID: 25952527.

14. Butterfield GE, Gates J, Fleming S, Brooks GA, Sutton JR, Reeves JT. Increased energy intake minimizes weight loss in men at high altitude. J Appl Physiol (1985) 1992;72(5):1741–8. doi:1G.1152/íappl.1992.72.5.1741 PMID: 1601781.

15. Karl JP, Cole RE, Berryman CE, Finlayson G, Radcliffe PN, Kominsky MT, et al. Appetite Suppression and Altered Food Preferences Coincide with Changes in Appetite-Mediating Hormones During Energy Deficit at High Altitude, But Are Not Affected by Protein Intake High Alt Med Biol. 2018. doi:10.1089/ham.2017.0155 PMID: 29431471.

16. Chen CY, Tsai YL, Kao CL, Lee SD, Wu MC, Mallikarjuna K, et al. Effect of mild intermittent hypoxia on glucose tolerance, muscle morphology and AMPK-PGC-1alpha signaling. Chin J Physiol. 2010;53(1):62–71. doi:10.4077/CJP.2010.AMK078

17. Lippl FJ, Neubauer S, Schipfer S, Lichter N, Tufman A, Otto B, et al. Hypobaric hypoxia causes body weight reduction in obese subjects. Obesity. 2010;18(4):675–81. doi:10.1038/oby.2009.509 PMID: 20134417.

18. Gonzalez-Muniesa P, Lopez-Pascual A, de Andres J, Lasa A, Portillo MP, Aros F, et al. Impact of intermittent hypoxia and exercise on blood pressure and metabolic features from obese subjects suffering sleep apnea-hypopnea syndrome J Physiol Biochem. 2015;71(3):589–99. doi:10.1007/s13105-015-0410-3 PMID: 25913417.

19. Balykin MV, Gening TP, Vinogradov SN. Morphological and functional changes in overweight persons under combined normobaric hypoxia and physical training Human physiology. 2004;30(2):184–191 doi:10.1023/b:hump.0000021647.73620 PMID: 15150977.

20. Netzer NC, Chytra R, Kupper T. Low intense physical exercise in normobaric hypoxia leads to more weight loss in obese people than low intense physical exercise in normobaric sham hypoxia Sleep Breath. 2008;12(2):129–34. doi:10.1007/s11325-007-0149-3 PMID: 18057976.

21. Gatterer H, Haacke S, Burtscher M, Faulhaber M, Melmer A, Ebenbichler C, et al. Normobaric Intermittent Hypoxia over 8 Months Does Not Reduce Body Weight and Metabolic Risk Factors– a Randomized, Single Blind, Placebo-Controlled Study in Normobaric Hypoxia and Normobaric Sham Hypoxia Obes Facts. 2015;8(3):200–9. doi:10.1159/000431157 PMID: 26008855.

22. Chandler PC, Viana JB, Oswald KD, Wauford PK, Boggiano MM. Feeding response to melanocortin agonist predicts preference for and obesity from a high-fat diet Physiol Behav. 2005;85(2):221–30. doi: 10.1016/j.physbeh.2005.04.011 PMID: 15893778.

23. Bagnol D, Alshamma HA, Behan D, Whelan K, Grottick AJ. Diet-induced models of obesity (DIO) in rodents Curr Protoc Neurosci. 2012;Chapter 9:Unit 9.38.1–13. doi:10.1002/0471142301.ns0938s59 PMID:22470151.

24. Bedford TG, Tipton CM, Wilson NC, Oppliger RA, Gisolfi CV. Maximum oxygen consumption of rats and its changes with various experimental procedures J Appl Physiol Respir Environ Exerc Physiol. 1979;47(6):1278. dio: 10.1152/jappal.1979.47.6.1278 PMID:536299.

25. Woods SC, Seeley RJ, Rushing PA, D’Alessio D, Tso P. A controlled high-fat diet induces an obese syndrome in rats J Nutr. 2003;133(4):1081–7.dio:10.1093/jn/133.4.1081 PMID:12672923.

26. Araújo EP, De Souza CT, Ueno M, Cintra DE, Bertolo MB, Carvalheira JB, et al. Infliximab restores glucose homeostasis in an animal model of diet-induced obesity and diabetes Endocrinology. 2007;148(12):5991.dio:10.1210/en.2007-0132 PMID:17761768.

27. Quintero P, Milagro FI, Campión J, Martínez JA. Impact of oxygen availability on body weight management Med Hypotheses. 2010;74(5):901–7. dio:10.1016/j.mehy.2009.10.022 PMID:19913361.

28. Urdampilleta A, Gonzalez-Muniesa P, Portillo MP, Martinez JA. Usefulness of combining intermittent hypoxia and physical exercise in the treatment of obesity J Physiol Biochem. 2012;68(2):289–304. doi: 10.1007/s13105-011-0115-1 PMID: 22045452.

29. Xi L, Chow CM, Kong X. Role of Tissue and Systemic Hypoxia in Obesity and Type 2 Diabetes J Diabetes Res. 2016;2016:1527852. doi:10.1155/2016/1527852 PMID: 27419143.

30. Gonzalez-Muniesa P, Quintero P, De Andres J, Martinez JA. Hypoxia: a consequence of obesity and also a tool to treat excessive weight loss. Sleep Breath. 2015;19(1):7–8. doi:10.1007/s11325-014-0972-2 PMID: 24807116.

31. Cerretelli P, Prampero PED, Howald H. Muscle Function Impairment in Humans Acclimatized to Chronic Hypoxia. Physiological Function in Special Environments.1989: 41–58 p.

32. Howald H, Hoppeler H. Performing at extreme altitude: muscle cellular and subcellular adaptations. Eur J Appl Physiol. 2003;90(3-4):360–4. doi:10.1007/s00421-003-0872-9 PMID: 12898262.

33. Hoppeler H, Kleinert E, Schlegel C, Claassen H, Howald H, Kayar SR, et al. Morphological adaptations of human skeletal muscle to chronic hypoxia. Int J Sports Med. 1990;11(S 1):S3–S9. doi:10.1055/s-2007-1024846 PMID:2323861.

34. Narici MV, Kayser B. Hypertrophic response of human skeletal muscle to strength training in hypoxia and normoxia. Eur J Appl Physiol Occup Physiol. 1995;70(3):213–9. PMID:7607195.

35. Ou LC, Leiter JC. Effects of exposure to a simulated altitude of 5500 m on energy metabolic pathways in rats. Respir Physiol Neurobiol. 2004;141(1):59–71. doi:10.1016/j.resp.2004.04.001 PMID: 15234676.

36. Qin L, Song Z, Wen SL, Jing R, Li C, Xiang Y, et al. Effect of intermittent hypoxia on leptin and leptin receptor expression in obesity mice. Sheng Li Xue Bao. 2007;59(3):351–6. PMID: 17579792.

37. Carreras A, Kayali F, Zhang J, Hirotsu C, Wang Y, Gozal D. Metabolic effects of intermittent hypoxia in mice: steady versus high-frequency applied hypoxia daily during the rest period. Am J Physiol Regul Integr Comp Physiol. 2012;303(7):R700–9. doi:10.1152/ajpregu.00258.2012 PMID: 22895743.

38. Ge RL, Wood H, Yang HH, Liu YN, Wang XJ, Babb T. The body weight loss during acute exposure to high-altitude hypoxia in sea level residents. Sheng Li Xue Bao. 2010;62(6):541–6. PMID: 21170501.

39. Mackenzie R, Maxwell N, Castle P, Elliott B, Brickley G, Watt P. Intermittent Exercise with and without Hypoxia Improves Insulin Sensitivity in Individuals with Type 2 Diabetes J Clin Endocrinol Metab. 2012;97(4):E546–E55. doi:10.1210/jc.2011-2829 PMID: 22278428.

40. Richalet JP, Larmignat P, Poitrine E, Letournel M, Canoui-Poitrine F. Physiological risk factors for severe high-altitude illness: a prospective cohort study Am J Respir Crit Care Med. 2012;185(2):192–8. doi: 10.1164/rccm.201108-1396GC PMID: 22071330.

41. Wee J, Climstein M. Hypoxic training: Clinical benefits on cardiometabolic risk factors J Sci Med Sport. 2015;18(1):56–61. doi:10.1016/i.isams.2013.10.247 PMID: 24268571.

42. Bailey DP, Smith LR, Chrismas BC, Taylor L, Stensel DJ, Deighton K, et al. Appetite and gut hormone responses to moderate-intensity continuous exercise versus high-intensity interval exercise, in normoxic and hypoxic conditions Appetite. 2015;89:237–45. doi:10.1016/j.appet.2015.02.019 PMID: 25700630.

43. Chen YC, Lee SD, Kuo CH, Ho LT. The effects of altitude training on the AMPK-related glucose transport pathway in the red skeletal muscle of both lean and obese Zucker rats High Alt Med Biol. 2011;12(4):371–8. doi:10.1089/ham.2010.1088 PMID: 22206563.

44. Lu YL, Jing W, Feng LS, Zhang L, Xu JF, You TJ, et al. Effects of hypoxic exercise training on microRNA expression and lipid metabolism in obese rat livers. J Zhejiang Univ Sci B. 2014;15(9):820–9. doi:10.1631/jzus.B1400052 PMID: 25183036.

45. He Z, Feng L, Zhang L, Lu Y, Xu J, Lucia A. Effects of hypoxic living and training on gene expression in an obese rat model. Med Sci Sports Exerc. 2012;44(6):1013–20. doi:10.1249/MSS.0b013e3182442d82 PMID: 22143106.

46. Roca-Rivada A, Castelao C, Senin LL, Landrove MO, Baltar J, Crujeiras AB, et al. FNDC5/Irisin Is Not Only a Myokine but Also an Adipokine Plos One. 2013;8(4):10. doi:10.1371/iournal.ŋone.0060563 PMID: 23593248.

47. Zhang Y, Li R, Meng Y, Li S, Donelan W, Zhao Y, et al. Irisin stimulates browning of white adipocytes through mitogen-activated protein kinase p38 MAP kinase and ERK MAP kinase signaling Diabetes. 2014;63(2):514–25. doi:10.2337/db13-1106 PMID: 24150604.

48. Polyzos SA, Anastasilakis AD, Efstathiadou ZA, Makras P, Perakakis N, Kountouras J, et al. Irisin in metabolic diseases Endocrine. 2018;59(2):260–74. doi:10.1007/s12020-017-1476-1 PMID: 29170905.

49. Choi YK, Kim MK, Bae KH, Seo HA, Jeong JY, Lee WK, et al. Serum irisin levels in new-onset type 2 diabetes Diabetes Res Clin Pract. 2013;100(1):96–101. doi:10.1016/j.diabres.2013.01.007 PMID: 23369227.

50. Liu JJ, Wong MDS, Toy WC, Tan CSH, Liu S, Ng XW, et al. Lower circulating irisin is associated with type 2 diabetes mellitus J Diabetes Complications. 2013;27(4):365–9. doi:10.1016/i.idiacomŋ.2013.03.002 PMID: 23619195.

51. Moreno-Navarrete JM, Ortega F, Serrano M, Guerra E, Pardo G, Tinahones F, et al. Irisin Is Expressed and Produced by Human Muscle and Adipose Tissue in Association With Obesity and Insulin Resistance J Clin Endocrinol Metab. 2013;98(4):E769–E78. doi:10.1210/jc.2012-2749 PMID: 23436919.

52. Stengel A, Hofmann T, Goebel-Stengel M, Elbelt U, Kobelt P, Klapp BF. Circulating levels of irisin in patients with anorexia nervosa and different stages of obesity – Correlation with body mass index. Peptides. 2013;39:125–30. doi:10.1016/j.peptides.2012.11.014 PMID: 23219488.

53. Kurdiova T, Balaz M, Vician M, Maderova D, Vlcek M, Valkovic L, et al. Effects of obesity, diabetes and exercise on Fndc5 gene expression and irisin release in human skeletal muscle and adipose tissue: in vivo and in vitro studies. J Physiol. 2014;592(5):1091–107. doi:10.1113/jphysiol.2013.264655 PMID: 24297848.

54. Pedersen BK, Febbraio MA. Muscles, exercise and obesity: skeletal muscle as a secretory organ. Nat Rev Endocrinol. 2012;8(8):457–65. doi:10.1038/nrendo.2012.49 PMID: 22473333.

55. Fatouros IG. Is irisin the new player in exercise-induced adaptations or not? A 2G17 update. Clinical chemistry and laboratory medicinel. 2018;56(4):525–48. doi:10.1.51.5/cclm-2017-0674 PMID: 29127759.

56. Fox J, Rioux BV, Goulet EDB, Johanssen NM, Swift DL, Bouchard DR, et al. Effect of an acute exercise bout on immediate post-exercise irisin concentration in adults: A meta-analysis. Scand J Med Sci Sports. 2018;28(1):16–28. doi:10.1111/sms.129G4 PMID: 28453881.

57. Bilski J, Mazur-Bialy AI, Brzozowski B, Magierowski M, Jasnos K, Krzysiek-Maczka G, et al. Moderate Exercise Training Attenuates the Severity of Experimental Rodent Colitis: The Importance of Crosstalk between Adipose Tissue and Skeletal Muscles. Mediat Inflamm. 2015:12. doi:10.1155/2015/605071 PMID: 25684862.

58. Andrade PA, Silveira BKS, Rodrigues AC, da Silva FMO, Rosa COB, Alfenas RCG. Effect of exercise on concentrations of irisin in overweight individuals: A systematic review. Sci Sports. 2018;33(2):8G–9. doi:0.1016/j.scispo.2017.11.002.

59. Norheim F, Langleite TM, Hjorth M, Holen T, Kielland A, Stadheim HK, et al. The effects of acute and chronic exercise on PGC-1 alpha, irisin and browning of subcutaneous adipose tissue in humans. FEBS J. 2014;281(3):739–49. doi:10.1111/febs.12619 PMID: 24237962.

60. Huh JY, Panagiotou G, Mougios V, Brinkoetter M, Vamvini MT, Schneider BE, et al. FNDC5 and irisin in humans: I. Predictors of circulating concentrations in serum and plasma and II. mRNA expression and circulating concentrations in response to weight loss and exercise. Metabolism. 2012;61(12):1725–38. doi:10.1016/j.metabol.2012.09.002 PMID: 23018146.

61. Scalzo RL, Peltonen GL, Giordano GR, Binns SE, Klochak AL, Paris HLR, et al. Regulators of Human White Adipose Browning: Evidence for Sympathetic Control and Sexual Dimorphic Responses to Sprint Interval Training PloS one. 2014;9(3). doi:10.1371/journal.pone.0090696 PMID: 24603718.

62. Sliwicka E, Cison T, Kasprzak Z, Nowak A, Pilaczynska-Szczesniak L. Serum irisin and myostatin levels after 2 weeks of high-altitude climbing PloS one. 2017;12(7). doi:10.1371/journal.pone.0181259 PMID: 28732027.

63. Becari C, Somers KR, Polonis K, Pfeifer MA, Prachi S. Acute Effects of Intermittent Hypoxia on Leptin-Mediated Increases in Adiponectin Expression Faseb Journal. 2016;30.

64. Fu C, Jiang L, Zhu F, Liu Z, Li W, Jiang H, et al. Chronic intermittent hypoxia leads to insulin resistance and impaired glucose tolerance through dysregulation of adipokines in non-obese rats Sleep Breath. 2015;19(4):1467–73. doi:10.1007/s11325-015-1144-8 PMID: 25724554.

65. Zhang X, Lam KSL, Ye H, Chung SK, Zhou M, Wang Y, et al. Adipose Tissue-specific Inhibition of Hypoxia-inducible Factor 1 alpha Induces Obesity and Glucose Intolerance by Impeding Energy Expenditure in Mice. J Biol Chem.2010;285(43):32869–77. doi:10.1074/jbc.M110.135509 PMID: 20716529.

66. De Matteis R, Lucertini F, Guescini M, Polidori E, Zeppa S, Stocchi V, et al. Exercise as a new physiological stimulus for brown adipose tissue activity Nutr Metab Cardiovasc Dis. 2013;23(6):582–90. doi:10.1016/j.numecd.2012.01.013 PMID: 22633794.

67. Chau MD, Gao J, Yang Q, Wu Z, Gromada J. Fibroblast growth factor 21 regulates energy metabolism by activating the AMPK-SIRT1-PGC-1alpha pathway. Proc Natl Acad Sci U S A. 2010;107(28):12553–8. doi:10.1073/pnas.1006962107 PMID: 20616029.

68. Cuevas-Ramos D, Aguilar-Salinas CA. Modulation of energy balance by fibroblast growth factor 21. Horm Mol Biol Clin Investig. 2016;30(1). pii:/j/hmbci.2017.30.issue-1/hmbci-2016-0023/hmbci-2016-0023.xml. doi:10.1515/hmbci-2016-0023 PMID: 27318658.

69. Shimizu I, Aprahamian T, Kikuchi R, Shimizu A, Papanicolaou KN, MacLauchlan S, et al. Vascular rarefaction mediates whitening of brown fat in obesity J Clin Invest. 2014;124(5):2099–112. doi:10.1172/jci71643 PMID: 24713652.

